# Dual-LLM Adversarial Framework for Information Extraction from Research Literature

**DOI:** 10.1101/2025.09.11.675507

**Authors:** Zhijing Li, Yunwen Yu, Wenhao Gu, Tiantian Zhu, Haohua Song, Wenbin Guo, Xiao Yang, Zexuan Zhu

## Abstract

Information Extraction (IE) is a fundamental task in Natural Language Processing (NLP) that aims to automatically identify relevant information from unstructured or semi-structured data. Information extraction from lengthy research literature, particularly in multi-omics studies, faces significant challenges due to their complex narratives and extensive context. To address this, we present a novel dual–LLM adversarial framework in which one large language model (LLM) performs the extraction and another provides iterative feedback to refine the results. This process systematically reduces errors, enhances consistency across heterogeneous data sources, and converges toward more accurate outputs. We evaluated our approach against manual and single-LLM extraction, using LLMs as evaluators. Experimental results show that our adversarial framework outperforms these baselines, highlighting its effectiveness for extracting structured information from lengthy scientific texts.

## 1 Introduction

The rapid growth of scientific publications has created an unprecedented wealth of knowledge, yet it also poses a significant challenge: how to efficiently identify, organize, and reuse critical information within massive volumes of text. Extracting such information from scientific literature is essential for accelerating discoveries, supporting reproducibility, and enabling downstream computational analyses.

Manual processing, however, is far too slow and resource-intensive to keep pace with the exponential expansion of the literature. Information Extraction (IE) techniques provide a potential solution by transforming unstructured text into structured, machine-readable knowledge. Nevertheless, traditional IE methods face several limitations: they often require large amounts of labeled training data, struggle with the long-range dependencies commonly found in scientific text, and lack the flexibility to generate outputs conforming to domain-specific structured schemas.

Recent advances in large language models (LLMs) offer new opportunities to overcome these barriers. LLMs excel at processing extended contexts and bring in domain knowledge from large-scale pretraining, making them promising for scientific IE tasks. Yet, obtaining high-quality results from LLMs in a single pass remains challenging. Issues such as incomplete coverage, inconsistency across outputs, and deviations from desired annotation formats continue to limit their reliability in practice.

To address these challenges, we propose a dual-LLM adversarial framework for scientific information extraction. In this framework, an Extractor LLM first produces structured outputs based on a domain-specific schema, while a Verifier LLM provides adversarial feedback to detect errors, omissions, and inconsistencies. Through iterative refinement, the two LLMs progressively improve the accuracy and consistency of the extracted information and its format alignment, enabling enhanced extractions without human intervention.

We evaluate our framework on multi-omics research literature, a domain characterized by complex data types and intricate semantic relationships. The results demonstrate that our approach significantly outperforms manual and single-LLM extraction, achieving higher accuracy and robustness in structured scientific information extraction. The innovations of our work are as follows.

- Adversarial Dual-LLM Framework – We introduce the first dual-LLM adversarial framework for scientific information extraction, where an Extractor LLM generates schema-guided extractions, and a Verifier LLM provides adversarial feedback. This closed-loop interaction enables automated self-correction without human intervention.
- IE Schema for Multi-Omics Studies: A customized schema that captures key elements-data, analyses and results-along with their interconnections.
- Multi-LLM-Based Scoring – A novel integration of multiple LLMs for scoring, providing a more comprehensive quality assessment.
- Empirical Validation with Multi-Omics Literature - Through extensive experiments, we show that our framework substantially outperforms manual and single-LLM extraction, demonstrating its effectiveness and scalability for processing complex scientific literature.

## 2 Related Work

IE has long been a central area of research in Natural Language Processing (NLP). Early efforts relied primarily on rule-based and statistical methods, such as BioIE Divoli and Attwood [2005] and PubTator Wei et al. [2024], to extract entities, relations, and events from structured and unstructured texts. While effective in specific domains, these approaches suffered from limited generalizability and required extensive domain expertise to manually craft rules.

The advent of deep learning transformed IE research. Sequence labeling models based on Bi-LSTM combined with Conditional Random Fields became the standard approach for tasks like named entity recognition and relation extraction Huang et al. [2015]Lample et al. [2016] Ma and Hovy [2016]. The introduction of transformer-based models, particularly BERT and its domain-specific variants such as SciBERT Beltagy et al. [2019] and BioBERT Lee et al. [2020], further advanced the state of the art in biomedical IE. These models improved contextual understanding and overall accuracy; however, extracting information from lengthy research papers remains a challenge due to complex discourse structure and long-range dependencies.

LLMs have further revolutionized IE by enabling longer context handling, integrating background knowledge, and performing extraction with minimal supervision Abu-Jeyyab et al. [2023]AI4Science and Quantum [2023]. GPT-based models and instruction-tuned variants have demonstrated remarkable performance in zero-shot and few-shot IE scenarios Hellberg [2023]. These models offer remarkable flexibility but often face challenges in consistency, factuality, and handling lengthy or multi-document contexts-common characteristics of multi-omics research literature. Several studies have attempted to address these limitations by using chain-of-thought prompting Wei et al. [2022]Diao et al. [2023] or integrating feedback mechanisms Yuksekgonul et al. [2025]. Automatic CoT generation, exemplified by Auto-CoT Zhang et al. [2022], matches the effectiveness of manually crafted prompts, while recent methods such as CoTGenius Cheng et al. [2024], which employs evolutionary strategies to generate more diverse, verifiable reasoning chains, leading to improved robustness and accuracy. Yet, comprehensive solutions for high-quality information extraction from lengthy scientific texts remain elusive.

Adversarial learning has also been successfully applied to various NLP tasks to improve model robustness and performance Li et al. [2017]Nie et al. [2019]. Generative adversarial networks (GANs) Goodfellow et al. [2014] have been adapted for text generation and classification tasks, demonstrating improved quality and diversity. Multi-agent systems represent another promising direction for complex NLP tasks. Recent work Du et al. [2023] has explored collaborative frameworks where multiple models work together to solve challenging problems.

Our study advances information extraction from scientific literature by introducing a dual-LLM adversarial framework. Unlike prior works that primarily benchmark LLM performance or rely on single-pass prompting Chen et al. [2025], our framework enables iterative refinement through an Extractor–Verifier interaction, thereby reducing errors and enhancing robustness. This design directly addresses the challenges of long-form scientific IE, where extended reasoning and domain precision are essential. Furthermore, while existing approaches often depend on string-matching metrics or manual evaluation Dagdelen et al. [2024], we propose a multi-LLM evaluator to capture contextual fidelity more effectively. Collectively, these contributions differentiate our work from prior efforts and underscore its novelty in tackling the pressing problem of reliable, high-quality information extraction from complex scientific texts.

## 3 Methodology

In this work, we introduce an adversarial dual–LLM framework designed to enhance the accuracy of information extraction from multi-omics literature. Traditional approaches are constrained by limited context and insufficient domain knowledge, while direct application of large language models fails to yield satisfactory results. Figure 1 shows the overall structure. Our method leverages two complementary LLMs: one dedicated to generating and refining extractions, and the other acting as a feedback model to evaluate their quality and provide targeted refinement guidance. Through iterative interactions, the framework progressively reduces errors and converges toward more accurate results. This section details the design principles, annotation schema, and iterative process underlying our proposed framework.

**Figure 1.**
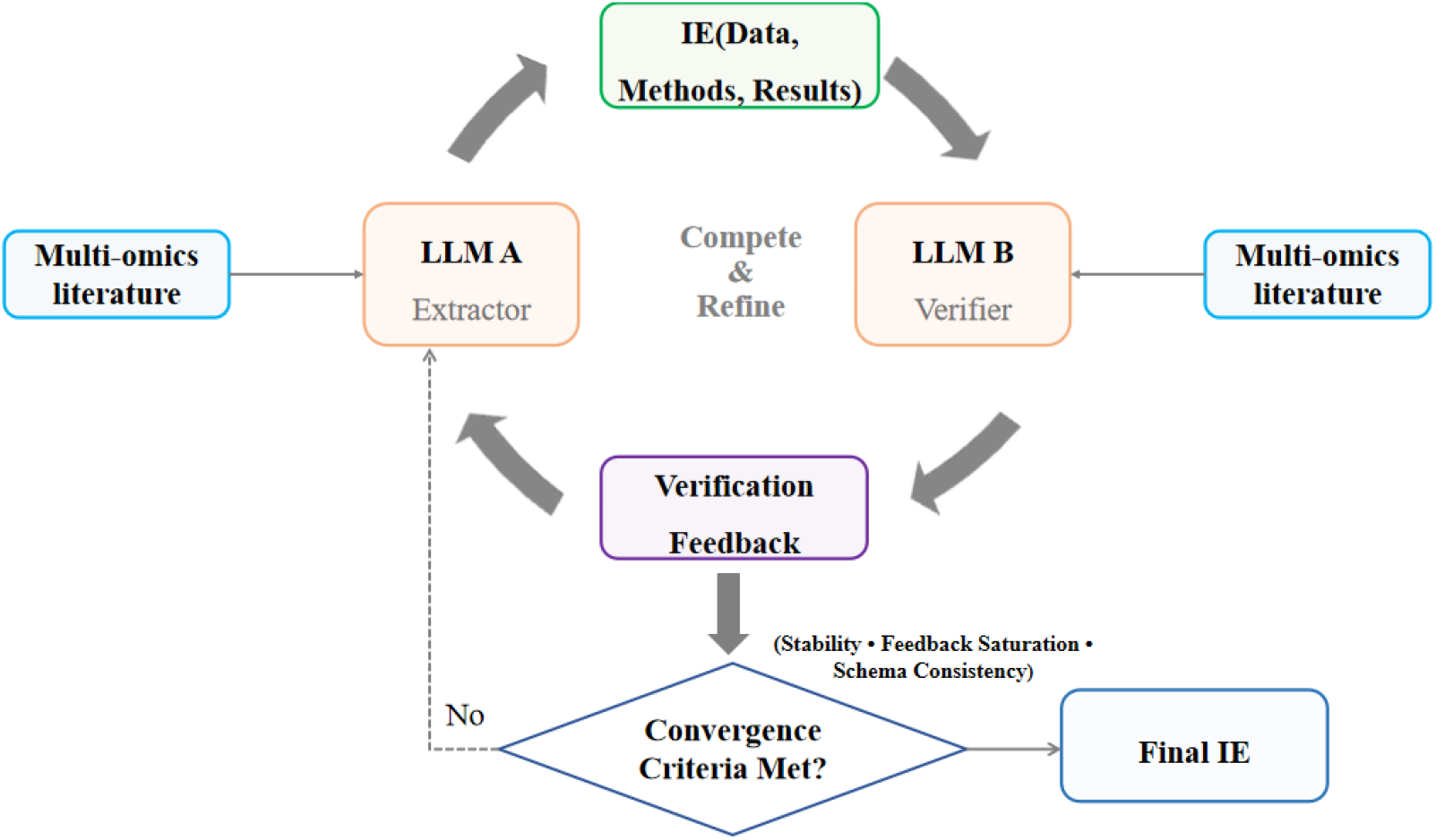
The overall structure of the dual-LLM adversarial frame

### 3.1 Extraction Rule Definition

In this paper, we have established extraction guidelines specifically for multi-omics research literature. The extraction guidelines consist of three primary components: data, analyses, and results. This tripartite structure is designed to capture the essential elements of multi-omics research. Specifically, the “data” component ensures that all used experimental data are accurately documented. The “analysis” section outlines the methodologies employed, enabling researchers to understand the analytical processes involved. Lastly, the “results” component emphasizes the importance of clearly presenting findings.

#### Definition of Data

In multi-omics research, data encompasses all forms of input information used in analyses and can be broadly classified into two primary categories:

1. Raw data: This includes unprocessed information collected directly from experimental platforms, such as:
  - Sequencing reads obtained through RNA-seq, DNA-seq, or other next-generation sequencing technologies.
  - Unprocessed mass spectrometry outputs used for analyzing protein composition.
  - Processed data: This refers to data that have been subjected to computational processing, including:
2. Normalized gene expression levels such as Transcripts.
  - Gene mutation profiles derived from variant calling pipelines that identify genetic variants.
  - Quantified protein abundance levels resulting from mass spectrometry or various proteomic techniques.

#### Definition of Analysis

Analysis involves a suite of computational and statistical methodologies that aims to uncover biological insights from experimental data. This process consists of several key stages:

1. Data preprocessing and transformation: This initial phase transforms raw experimental data-such as RNA-seq reads, genomic variant calls, or protein quantification measurements—into interpretable formats. The processed outputs, including quantifiable gene expression levels, detailed mutation profiles, and protein abundance metrics, form the foundation for subsequent analyses.
2. Bioinformatics analysis methods: This segment employs various techniques to elucidate the biological significance of the processed data. Key methods include:
  - Differential gene expression analysis: Identifying genes with statistically significant changes in expression levels under varying conditions.
  - Gene set enrichment analysis (GSEA): Detecting pathways or gene sets that are enriched in specific biological contexts, providing insights into relevant biological processes.
  - Statistical modeling: Applying statistical techniques to interpret biological implications and relationships within the data.
3. Machine Learning Analysis Methods: This component leverages machine learning algorithms to model and predict biological outcomes. Techniques employed in this phase include:
  - Classification analysis: Building models to classify samples into distinct groups, such as disease versus control.
  - Regression analysis: Modeling continuous outcomes, such as drug response level.
  - Survival analysis: Predicting time-to-event outcomes, such as patient survival.

#### Definition of Result

The results represent the outputs generated from the analyses, organized into two main categories:

1. Bioinformatics analysis outputs: This category includes the identification of differential biological entities, such as genes, cells, and pathways, along with statistical and computational metrics that quantify their significance. Metrics such as p-value, fold change, and enrichment score collectively indicate the magnitude and relevance of observed biological differences or relationships.
2. Machine learning performance metrics and features: This section pertains to the evaluation of machine learning models applied to predict biological outcomes. Key performance indicators, including Area Under the Curve (AUC), F1 score, sensitivity, and specificity, are reported to assess model effectiveness. Additionally, the features used in modeling, such as genes, proteins, and other biomarkers-are provided to facilitate interpretation and understanding of the underlying biological processes.

### 3.2 IE Format

The IE format is designed to systematically capture the essential elements of multi-omics research—data, analyses, and results—by explicitly linking each dataset to the specific analyses it informs and the results they generate. This ensures these components are treated as interconnected rather than isolated. Below is the detailed description of each component:

**Data Format:** <data_id, omics_type, data_link, data_format, data_source>

- data_id: Unique identifier for the data.
- omics_type: Type of omics data, including: transcriptomics, genomics, proteomics, or other applicable categories.
- data_link (optional): URL or reference link for online dataset access.
- data_format: File format specification, including fastq, bam, maf, vcf, txt, csv, tsv, or other applicable formats.
- data_source (optional): The origin or repository from which the data was obtained.

~~~
“data”: [
{
“id”: “<data_id>“,
“omics_type”: “<omics_type>“,
 “link”: “<data_link>“,
 “format”: “<data_format>“,
“source”: “<data\_source>“
}
]
~~~

**Analysis Format:** <analysis_id, analysis_type, analysis_data, training_set, test_set, label>

- analysis_id: Unique identifier for the conducted analysis.
- analysis_type: Type of performed analysis, including: RNA-seq, variant calling, protein quantification, differential analysis, single cell clustering, classification analysis, regression analysis, survival analysis, or other applicable methods.
- analysis_data: List of data_id entries utilized in the analysis.
- training_set (optional): List of data_id entries comprising the training dataset for machine learning analyses.
- test_set (optional): List of data_id entries comprising the test set for machine learning analyses.
- label (optional): Dictionary containing label information utilized in the analysis.

~~~
“analyses”: [
{
“id”: “<analysis_id>“,
“analysis_type”: “<analysis_type>“,
“analysis_data”: [“<data_id_1>“, “<data_id_2>“],
“training_set”: None,
“test_set”: None,
“label”: {
“<label_name>“: [“<group_name_1>“, “<group_name_2>“]
}
}
]
~~~

**Result Format:** <analysis_id, metric, performance, biomarkers>

- analysis_id: Identifier for the analysis corresponding to which the results were obtained.
- metric: Evaluation metric for the results, including: p-value, fold change, AUC, F1 score, or other applicable measures.
- performance: Overall performance assessment or a list of performance metrics for each biomarker, measured using the above metric.
- biomarkers: (List) List of the identified biomarkers (e.g., genes, proteins) from the analysis.

~~~
“results”: [
{
“corresponding_analysis_id”: “<analysis_id>“,
 “metrics”: “<metric>“,
“performance”: “<performance>“,
“biomarkers”: [“<biomarker_1>“, “<biomarker_2>“]
}
]
~~~

### 3.3 Structured Prompting

We designed a structured prompting approach to enhance extraction accuracy by defining clear formats and requirements, reducing errors caused by ambiguity. This method breaks down complex tasks into distinct parts, allowing the model to grasp key points and understand instructions more quickly. Information is divided into sections such as ‘data,’ ‘analyses,’ and ‘results,’ aligning with the annotation definition. Structured prompting ensures that the model captures critical information, providing a comprehensive understanding of the literature.

### 3.4 Dual-LLM Adversarial Framework

We introduce a dual-LLM adversarial framework for information extraction from multi-omics research literature, designed to enhance content accuracy and format consistency across heterogeneous biomedical sources. The framework consists of two complementary components: an Extractor LLM, which extracts information according to a domain-specific multi-omics schema, and a Verifier LLM, which iteratively evaluates the extractions to detect errors and missing information. Through an adversarial feedback loop, the Verifier guides the Extractor to refine outputs progressively, yielding high-fidelity results that capture complex information from multi-omics research.

### 3.5 Adversarial Iteration

The adversarial iteration process is central to the proposed framework, enabling systematic refinement of extractions through structured interaction between two complementary large language models. Initially, the Extractor LLM generates preliminary outputs directly from the literature. However, due to the inherent ambiguity and complexity of multi-omics research, the initial outputs are often susceptible to content errors, redundancy, and the omission of subtle contextual cues.

To address this limitation, the Verifier LLM generates detailed feedback in the subsequent step. It critically evaluates the Extractor’s output by identifying inconsistencies with the domain schema, highlighting errors, and detecting gaps where essential information is missing. Beyond error detection, it provides targeted refinements, offering explicit guidance for improving extracted data, analyses and results, clarifying their interconnections, and ensuring alignment with domain-specific conventions.

The Extractor then incorporates this feedback to refine its outputs. Through this iterative refinement, the adversarial loop systematically reduces both content and structural errors, while enhancing the robustness of extracted information across heterogeneous biomedical sources.

### 3.6 Evaluation

To assess the extracted results automatically, we adopt a multi-LLM–based evaluation protocol. Specifically, multiple LLMs are provided with both the extracted outputs and the ground truth, are instructed to assign two types of scores: a content accuracy score (0-100) and a format score (0-100), reflecting the consistency between the results and the ground truth in terms of both content and presentation. The final evaluation score is calculated as the average of all evaluators’ combined content and format scores.

When selecting and configuring LLMs for scoring, the following criteria are applied:

- Correct results should receive high scores.
- Incorrect results should receive low scores.
- Multiple evaluations of the same result should yield consistent scores.
- Correct results presented in different orders should receive the same score.

## 4 Experiments

We downloaded 42 multi-omics research papers on different diseases, each ranging from 37 to 48 pages to ensure diversity and representativeness. Each paper was annotated by two independent bioinformatics researchers according to the standards we set and verified by a third reviewer. These annotations serve as the ground truth. Half of them were used as a validation set for refining prompts and tuning parameters such as the choice of LLMs, while the remaining papers were reserved as a test set for performance evaluation and comparison.

### 4.1 LLM Evaluator

Qwen3-32B, Llama3.3-49B, GPT-4.1, GPT-4o, and GPT-o3—representing both open-source and commercial families of state-of-the-art LLMs—were initially selected as candidate evaluators. These candidates were tested against three ground-truth variants—original, corrupted, and reordered—to assess their adherence to the four evaluation criteria described above.

As shown in Table 1, most candidate models performed well in distinguishing correct from incorrect outputs, reliably assigning high scores to correct annotations and low scores to corrupted ones. However, further stress-testing revealed key limitations. In particular, Llama3.3-49B did not perform satisfactorily under the Order Invariance setting, producing inconsistent scores when the same set of annotations was presented in different orders. This indicates that Llama3.3-49B does not fully meet the robustness requirements for serving as a reliable evaluator.

**Table 1:** Performance of different LLM evaluators under four selection criteria

| Evaluator | Correct Results | Incorrect Results | Consistency | Order Invariance |
| --- | --- | --- | --- | --- |
| Qwen3-32B | ✓ | ✓ | ✓ | ✓ |
| Llama3.3-49B | ✓ | ✓ | ✓ | × |
| GPT series (GPT-4.1, GPT-4o, GPT-o3) | ✓ | ✓ | × | × |

Similarly, the GPT family showed instability across repeated evaluations: despite identical seeds and temperatures, the same annotation often received noticeably different scores, compromising reproducibility.

In contrast, Qwen3-32B consistently satisfied all evaluation criteria, demonstrating both stability and robustness across original, corrupted, and reordered conditions. Therefore, Qwen3-32B was ultimately chosen as the sole evaluator for our experiments.

### 4.2 Single-LLM Extraction (Non-Adversarial)

We first establish a baseline by evaluating single LLMs in a one-pass information extraction setting on the validation set, where each model is prompted with schema-specific instructions without iterative refinement. The results are summarized in Table 2. Qwen3-32B achieved the best overall performance with an evaluation score of 52.5, closely followed by Qwen2.5-32B (51.18), demonstrating that larger Qwen models are relatively more robust in information extraction from lengthy text. Llama3-49B and GPT-4o reached moderate scores around 47, while Gemma3-27B and Mistral-Small3.2-24B performed less competitively, with evaluation scores dropping to 46.22 and 38.82, respectively. These findings highlight that the effectiveness of single-pass prompting is highly influenced by the choice of LLM, with considerable performance gaps observed across different models. Moreover, even the strongest single-LLM baselines show limitations, motivating the exploration of dual-LLM adversarial frameworks for more reliable information extraction.

**Table 2:** Performance of different single LLMs on the validation set

| LLM | Evaluation Score |
| --- | --- |
| Qwen3-32B | 65.06 |
| Qwen2.5-32B | 60.43 |
| Llama3-49B | 58.06 |
| Gemma3-27B | 59.21 |
| Mistral-Small3.2-24B | 47.68 |
| GPT-4o | 57.62 |

### 4.3 Dual-LLM Adversarial (Proposed)

Our method introduces an adversarial loop between an Extractor LLM and a Verifier LLM. The Extractor produces initial schema-based outputs, while the Verifier identifies errors, inconsistencies, and omissions. Table 3 reports the performance of different Extractor–Verifier pairings over two iterations on the validation set.

**Table 3:** Performance of different Extractor–Verifier pairings on the validation set

| Extractor LLM | Verifier LLM | Evaluation Score |
| --- | --- | --- |
| Qwen3-32B-no-thinking | Qwen3-32B-no-thinking | 70.81 |
| Qwen3-32B-no-thinking | Qwen3-32B-thinking | 66.17 |
| Qwen3-32B-thinking | Qwen3-32B-thinking | 67.17 |
| Qwen3-32B-thinking | Qwen3-32B-no-thinking | 66.49 |
| Llama-3.3-nemotron-49b-super-v1-no-thinking | Llama-3.3-nemotron-49b-super-v1-no-thinking | 62.24 |
| Llama-3.3-nemotron-49b-super-v1-no-thinking | Llama-3.3-nemotron-49b-super-v1-thinking | 61.02 |
| Llama-3.3-nemotron-49b-super-v1-thinking | Llama-3.3-nemotron-49b-super-v1-thinking | 62.97 |
| Llama-3.3-nemotron-49b-super-v1-thinking | Llama-3.3-nemotron-49b-super-v1-no-thinking | 65.16 |
| GPT-4o | GPT-4o | 69.98 |

Within the Qwen3-32B family, evaluation scores are generally high, ranging from 66.17 to 70.81. The best result (70.81) is obtained when both the Extractor and Verifier are configured without “thinking,” whereas introducing a “thinking” Verifier tends to reduce performance. For the Llama-3.3-nemotron models, performance is lower overall, with scores between 61.02 and 65.16, but still shows moderate variation depending on the pairing. GPT-4o, when used as both Extractor and Verifier, also performs strongly, achieving 69.98, comparable to the best Qwen results.

These findings demonstrate that the choice of Extractor–Verifier combination has a measurable impact on information extraction quality. In particular, while Qwen3-32B models exhibit the highest overall scores, Llama-based models lag behind, and GPT-4o achieves competitive performance. This provides clear evidence that pairing strategy plays a critical role in optimizing the dual-LLM adversarial framework.

Figures 2 present the score distributions when using Qwen3-32B-no-thinking as both Extractor and Verifier, evaluated by Qwen3-32B across four dimensions: data, analysis, result, and average score. The evaluator reveals a consistent pattern: the highest scores appear in the data dimension, moderate scores in analysis, and the lowest scores in results. This trend is intuitive—multi-omics research papers typically present the data section in a clear and structured manner, whereas results are more complex and dispersed across text, tables, and figures, with analyses falling in between.

**Figure 2.**
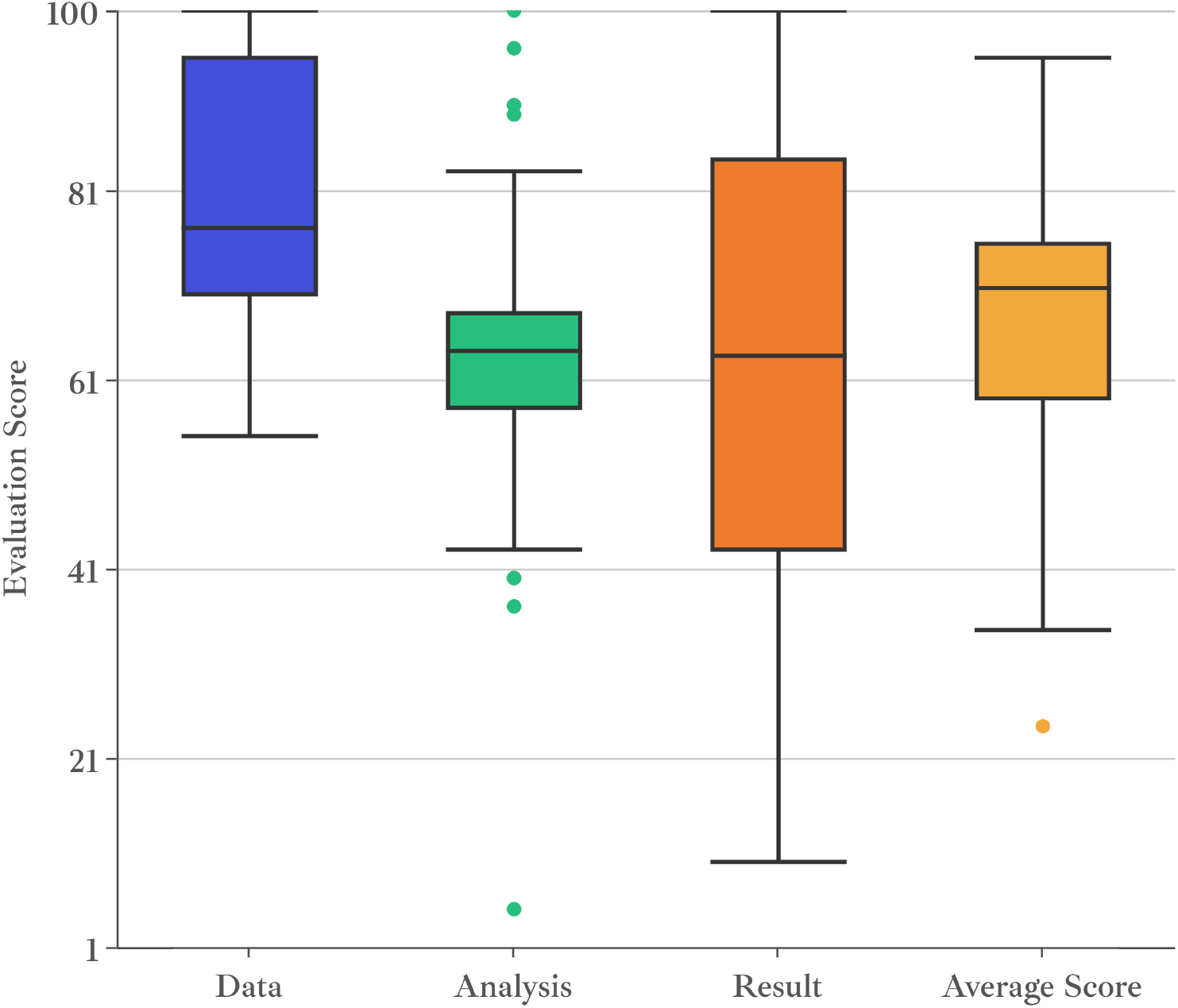
Score distribution of Qwen3-32B evaluator

#### Effect of Iteration Rounds

We further examined the impact of the number of adversarial rounds on performance, using Owen3-32B-thinking as the Extractor and Qwen3-32B-no-thinking as the Verifier. As shown in Figure 3, the system achieves its best results at the second iteration, where both content accuracy and structural consistency reach their peak. This indicates that the Verifier LLM provides the most effective corrections during the early refinement stage, resolving major errors and structural inconsistencies in just a few rounds. Beyond the second iteration, however, improvements diminish, and in some cases performance slightly decreases due to overcorrection or unnecessary modifications. After approximately ten iterations, results converge, with only marginal gains observed in subsequent rounds. This suggests that the adversarial framework effectively resolves most content errors, omissions, and structural inconsistencies within the first ten iterations, while further refinement primarily introduces negligible adjustments. Figure 4 illustrates the score changes of each paper before and after adversarial refinement.

**Figure 3.**
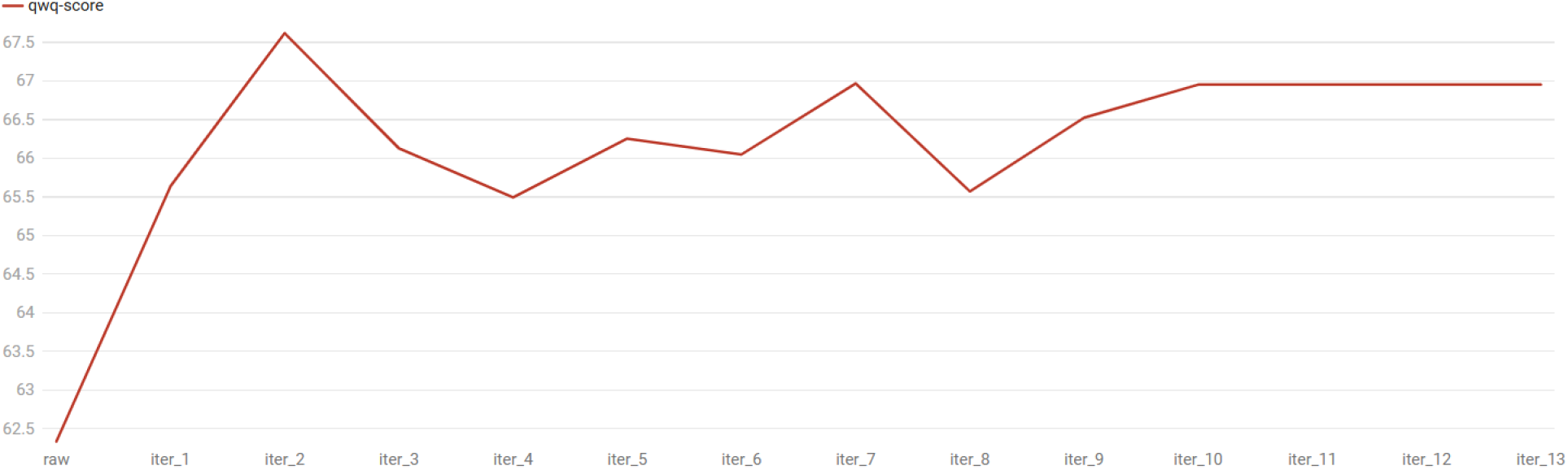
Evaluation score dynamics across adversarial refinement rounds

**Figure 4.**
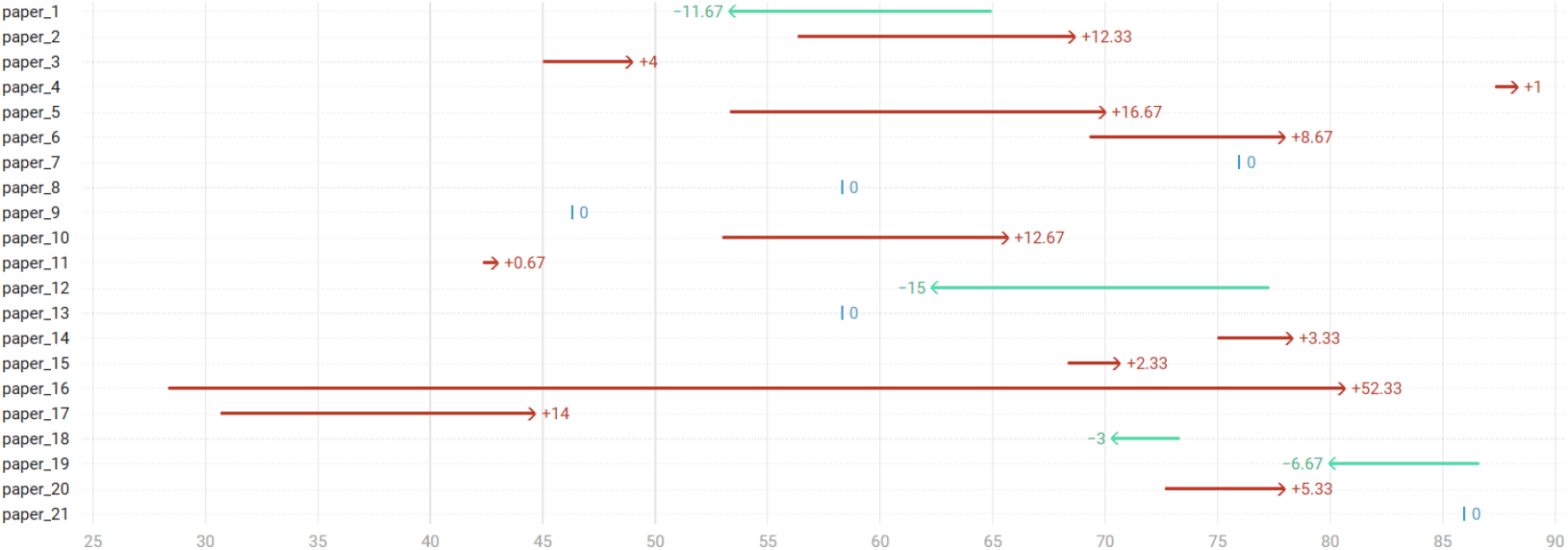
Evaluation score variations of each paper before and after adversarial refinement

Consequently, two iterations represent the optimal balance between performance and efficiency, delivering better results without incurring additional computational cost or risking instability from excessive feedback.

### 4.4 Comparative Experiments

To assess the effectiveness of our framework, we compare it against both manual single-LLM extraction on the test set of 21 papers. Based on the validation set experiments, we selected Qwen3-32B as the representative single-LLM baseline due to its consistently superior performance among tested models. For the adversarial setting, we adopted the Qwen3-32B-no-thinking - Qwen3-32B-no-thinking pairing, which yielded the strongest results during validation, thereby serving as the dual-LLM configuration for comparison.

#### Manual Extraction

Domain experts in bioinformatics manually extracted a subset of the literature according to the multi-omics IE schema. To establish a manual performance benchmark, we recruited seven experts with bioinformatics backgrounds to manually annotate benchmark literature, with each expert randomly assigned three different papers. As a result, each paper received IE from a single experts.

Table 4 reports the comparative performance of manual extraction, the single-LLM baseline, and the dual-LLM adversarial framework on the held-out test set. Manual extraction yields the lowest overall score, highlighting the difficulty of consistently capturing complete and well-structured information from lengthy multi-omics texts. A single LLM substantially outperforms manual extraction (56.76 vs. 44.89), demonstrating the stability of large language models in schema-guided extraction. Notably, the dual-LLM adversarial framework achieves the best overall performance, exceeding the single-LLM baseline by approximately 6.5 points.

**Table 4:** Evaluation score comparison between manual extraction, single-LLM and dual-LLM adversarial on the test set

|  | Manual Extraction | Single-LLM | Dual-LLM |
| --- | --- | --- | --- |
| Data | 57.43 | 76.29 | 75.1 |
| Analyses | 42.1 | 47.05 | 64.9 |
| Results | 35.13 | 46.93 | 49.8 |
| Average | 44.89 | 56.76 | 63.27 |

A breakdown across dimensions reveals that the improvement is not uniform. In the Data dimension, both the single and dual models achieve high scores (76.29 and 75.1, respectively), suggesting that factual data extraction is relatively straightforward and leaves limited room for adversarial refinement. In contrast, the Analyses dimension exhibits the largest margin of improvement: the dual model reaches 64.9, significantly higher than both the single model and manual extraction. This indicates that the adversarial interaction is particularly effective in capturing methodological details and logical reasoning steps, which are prone to errors and omissions in single-pass extraction. For the Results dimension, the dual framework delivers only modest gains over the single model (49.8 vs. 46.93), reflecting the inherent complexity of results sections, where information is often fragmented across prose, tables, and figures.

Overall, these findings confirm that the proposed adversarial framework provides substantial benefits over both manual and single-model baselines, especially for analysis-oriented content, while also exposing persistent challenges in extracting highly dispersed and structurally complex result information..

## 5 Discussion

In this work, we develop a novel dual-LLM Adversarial Framework for information extraction, applied to multi-omics research literature to capture experimental data, analytical methods, results, and their complex interconnections. To evaluate extraction quality, we employ Qwen3-32B as the sole evaluator, which scores outputs on both content accuracy and format consistency against ground truth. The framework is benchmarked on 21 multi-omics papers, comparing its performance with manual and single-LLM extraction. Through iterative adversarial interaction, the framework progressively refines its outputs, achieving superior performance relative to both manual and single-LLM approaches. These results highlight the framework’s effectiveness for extracting structured information from lengthy scientific texts.

An analysis of performance trends across adversarial refinement rounds reveals marked improvements during the first two iterations, followed by a slight decline and eventual convergence within ten iterations. This decline can be attributed to overcorrection and unnecessary modifications, suggesting that the adversarial process requires careful calibration to avoid diminishing returns. This finding provides valuable insights for optimizing iterative refinement strategies in dual-LLM architectures.

Despite outperforming manual and single-LLM extraction, our framework exhibits a measurable gap compared to ground truth, indicating that the Verifier LLM cannot fully identify all errors, omissions, and format inconsistencies produced by the Extractor LLM. This limitation suggests opportunities for enhancing the verification component through more sophisticated error detection mechanisms.

Another limitation is the relatively small scale of our evaluation, which involved only 21 multi-omics papers. This constrained sample size may limit the generalizability of our findings. Future work should assess the framework on larger corpora encompassing lengthier texts and diverse research domains.

Finally, while this study employs Qwen3-32B as the sole evaluator, relying on a single model may introduce bias or blind spots. In particular, in the test set experiments we used Qwen3-32B for the single-LLM baseline and the Qwen3-32B-no-thinking - Qwen3-32B-no-thinking configuration for the dual-LLM setting. Since Qwen3-32B also serves as the evaluation model, it is possible that the evaluator exhibits a preference toward outputs generated by models from the same family, leading to systematically higher scores. This potential bias underscores the importance of diversifying evaluators. Future work will therefore extend the evaluation strategy to incorporate multiple heterogeneous LLM-based evaluators, complementing the limitations of any single model and enhancing the robustness and fairness of performance assessment.

## 6 Conclusion

In this study, we present the first attempt to extract multi-omics information in a structured data–analysis–result manner, enabling a more systematic representation of scientific knowledge. Our experiments demonstrate that the adversarial framework, which combines extractor and verifier models, consistently outperforms both single-model and manual labeling, achieving higher accuracy and robustness. Importantly, this framework effectively mitigates inconsistency and schema deviation that often arise in single-pass LLM extraction, thereby improving reliability when processing complex multi-section scientific texts. Furthermore, we introduce an evaluation paradigm based on LLM evaluator, allowing for more nuanced and context-aware assessments beyond traditional string-matching metrics. Together, these findings highlight the promise of adversarial LLM frameworks as a scalable and accurate solution for IE from biomedical literature.

